# The role of shape and local features in human object recognition and learning

**DOI:** 10.64898/2026.11.12.765986

**Authors:** Allan R. Schneider, Olivia Maltz, Vladislav Ayzenberg

## Abstract

An object’s shape is an important visual cue, critical for both perception and cognition. However, the current best models of human vision, Deep Neural Networks (DNNs), do not prioritize shape, but instead, they rely on texture-like local visual features. This raises the question: Could shape be less important for human perception than previously believed? In seven experiments with both adults (*N* = 300) and 5-year-old children (*N* = 100), we tested the role of shape and local features in visual object recognition and learning. We found that humans could identify common objects as accurately using local visual features as they could using shape. This finding suggests that, like DNNs, local features may be sufficient for everyday visual recognition and that shape may not be necessary. However, we found that shape played a crucial role in novel object learning, allowing participants to even accomplish one-shot object categorization. Together, these findings force us to reconsider the role of shape in visual recognition and suggest that DNNs, despite not using shape, may nevertheless capture an important aspect of human visual processing.

**Significance:** Computational models have long sought to emulate the visual recognition and learning abilities of humans. Yet, the current best models neglect a crucial aspect of human vision: shape perception. Instead, models overwhelmingly rely on texture-like local features. Here, we investigated whether local features may play a more important role in human visual recognition than previously believed. We found that adults and children could accurately identify common objects using only local features, and shape was not needed. However, shape played an especially important role when participants needed to learn new object categories. These findings shift our current understanding of the role of shape and local features in visual perception, and reconcile longstanding debates regarding the alignment of machines to humans.

## Introduction

Object shape is often considered one of the most important visual cues underlying perception and cognition. Humans accurately recognize objects by their shape in the absence of other cues (e.g., line drawings) (1, 2), and preferentially rely on shape information across variations in viewpoint, category exemplar, and conflicting color and texture information (3–6). Shape perception also arises early in development, such that even newborn infants exhibit shape constancy (7), and by 6-months-of-age, use shape to rapidly learn new categories (8–11). Moreover, shape is the principal cue underlying children’s first-learned words (4), and is the perceptual scaffold of most basic-level categories (12)—the categorization level privileged for recognition speed, naming, and memory (13, 14). More broadly, by defining the boundaries of objects, shape information helps subdivide the visual world into meaningful perceptual units that become the targets of attention and action (15). Thus, shape not only carves up the visual world, but also underpins and grounds our conceptual ontology—our sense of what kinds of things exist as discrete, nameable objects.

Despite the importance of shape in human vision, the current best computational models of vision, Deep Neural Networks (DNNs), do not seem to use shape. In contrast to humans, the visual recognition performance of DNNs drops dramatically when tested using shape stimuli (e.g., line drawings) (16–18), and even with specialized training meant to emphasize object shape (18, 19), their recognition of object shape is often worse than that of infants and young children (11, 20). Instead, DNNs appear to prioritize local image features, such as the texture or local object parts (16, 18, 19, 21–23). For instance, when presented with objects whose shape and texture correspond to different categories, conventional DNNs will classify them on the basis of texture rather than shape (16, 18, 19). This local feature bias is sufficiently strong that even small perturbations to an object image, that may be imperceptible to humans, are sufficient to change the DNN’s predicted label for that object (16, 24, 25). DNN’s difficulties in representing shape appears to be rooted in their inability to represent the spatial relations between object parts (26–28) which forms the basis of robust shape representations, allowing humans to prioritize global form over local contours or features (3, 29–31). For instance, DNNs’ accuracy remains high when global shape information, but not local features, is disrupted, such as by spatially rearranging the position of object parts (16, 21, 23, 32–34).

Nevertheless, without representing shape information, DNNs exhibit remarkable performance on a range of visual tasks and provide the current best computational description of primate visual behavior and neural responses (35, 36). DNNs often match or surpass humans on tasks ranging from naturalistic image recognition (37), to medical image analysis (38, 39). Moreover, human object recognition performance and similarity judgments are consistently well predicted and matched by DNNs across a range of tasks (40–42). Additionally, like the primate visual system, DNNs develop hierarchical representations, with visual features becoming increasingly more complex as information progresses towards higher layers (43–46). And, indeed, the internal responses of DNN units show a high degree of quantitative alignment with the neural representations of the primate visual system, both at the single-neuron (47, 48) and population-activity levels of analysis (36, 43, 44, 49, 50).

How might DNNs achieve such performance and biological similarity without representing a seemingly fundamental cue such as shape? One possibility is that DNNs are not in fact good models of human vision (51, 52). For instance, they may achieve strong performance using different strategies than humans (53, 54) and their fit to biological data may arise from flexibility in the fitting process (50). However, an often-overlooked alternative possibility is that DNNs *are* in fact capturing an important aspect of human visual perception. Specifically, the success and fit of DNNs to human vision may suggest that shape information is less important than historically believed. In this view, local visual features may be sufficient for humans to accomplish many of the same visual tasks that DNNs succeed at, and thereby also explain DNNs’ strong match to the human visual system.

In the current study, we tested the importance and functional role of shape and local features in human visual perception. In the first set of experiments, we tested whether local visual features, in the absence of shape, were sufficient for humans to recognize familiar, everyday objects. To this end, adults were presented with either object silhouettes, which preserve global shape information but disrupt local features, or spatially scrambled object images, which preserve local features but disrupt global shape information (Fig. 1A). Then, in a second set of experiments, we explored the functional role of shape in object recognition and specifically tested whether it may support rapid category learning. Here, participants were presented with musical instruments from different cultures—objects that are novel to participants but have a real-world category structure—and we examined the extent to which shape and local features support one-shot category learning. Finally, we examined the developmental stability of these abilities by testing the extent to which 5-year-old children are similarly capable of recognizing and learning objects using shape or local features.

**Figure 1.**
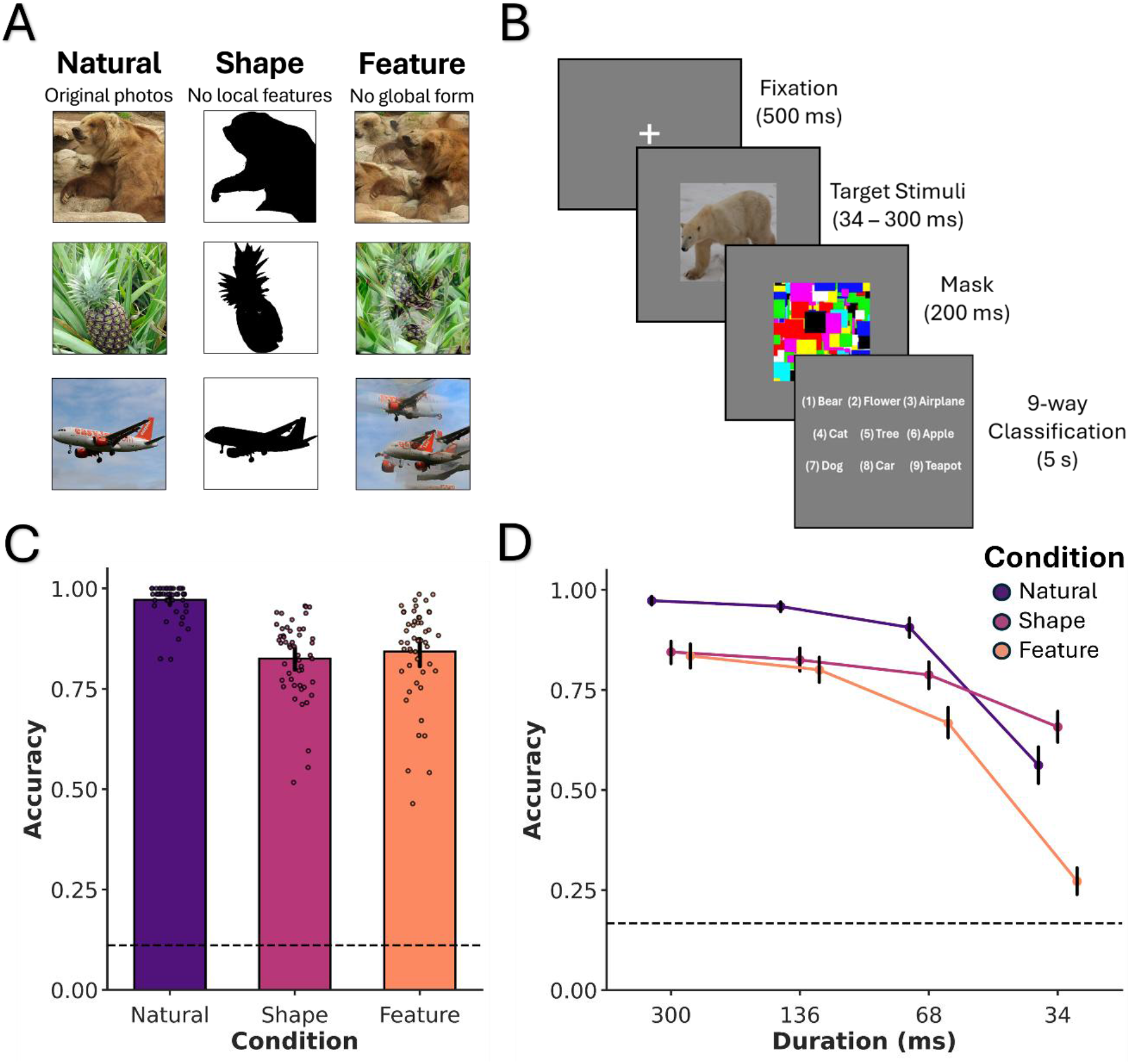
Familiar object recognition conditions, task, and results. (A) Example stimuli from the natural, shape, and feature conditions. Shape stimuli were generated using naturalistic objects that were transformed into 2D silhouettes, which preserved the global form but removed the local features. Feature stimuli were naturalistic objects that were spatially scrambled, which preserved the local features but disrupted the global form of the object. (B) On each trial, participants were required to identify a rapidly presented object image at either a consistent duration (300 ms) or a variable duration (300, 136, 68, 34 ms), from among a set of labels (consistent duration: 9 labels; variable duration: 6 labels). (C-D) Participants performance for natural, shape, and feature conditions in the (C) consistent and (D) variable duration experiments. (C-D) Participant performance was above chance in all conditions, and all stimulus durations. Individual points correspond to each participant’s mean accuracy on that condition. Error bars correspond to the upper and lower bounds of 95% confidence intervals. Dashed lines correspond to chance performance (consistent duration: 0.11; variable duration: 0.16).

To foreshadow our results, in both adults and children we found that local features were sufficient for everyday visual object recognition, such that recognition accuracy for familiar objects was as high using local features as when they used shape. However, shape was the dominant cue when learning new object categories, allowing participants to form new categories from just one exemplar. Together, these computationally inspired experiments provide new insights into the mechanisms underlying human visual perception and learning across development. Moreover, these data reconcile long-standing contradictory findings regarding the similarity of DNNs and the human visual system.

## Results

### Local features are sufficient for familiar, naturalistic object recognition

We first examined the extent to which local features were sufficient to categorize familiar, everyday objects. Adult participants (*n* = 50) were shown a rapidly presented object and asked to categorize the object from list of labels (Fig. 1B). The labels consisted of object classes both within and outside the same subordinate category as the target object, namely, animate (e.g., bear), inanimate natural (e.g., pineapple), and inanimate artificial (e.g., airplane). Importantly, the stimuli were presented in one of three conditions: (1) a ‘natural condition’ wherein the image is shown in its original undisrupted state, (2) ‘shape condition’ wherein the local features are disrupted by presenting only the object silhouette, but global shape information is preserved, and a (3) ‘feature condition’ wherein the local features are preserved but global shape information is disrupted via spatially scrambling the image (Fig. 1A).

One-sample *t*-test comparisons to chance (0.11), revealed participants exhibited above chance performance for natural (*M* = .97), shape (*M* = .83), and local feature (*M* = .84) conditions (*ps* < .001, *ds* > 6.17, BF₁₀ > 100; Fig. 1C), providing initial evidence that local features are sufficient for object recognition. A repeated-measures ANOVA with stimulus condition as the within-subjects factor revealed a main effect of condition on performance (*F*(2,98) = 68.09, *p* < .001, *n^2^_p_* = .58, BF_10_ > 100), with post-hoc multiple comparisons (Holm-Bonferroni corrected) showing that, unsurprisingly, performance on the natural condition was significantly higher than both shape and feature conditions (*ps* < .001, *ds* > 1.40, BF_10_ > 100). Interestingly, however, there was no difference between accuracy on shape and local feature conditions (*t*(49) = 1.26, *p* = .214, *d* = 0.19, BF_10_ = 0.32), and, in fact, accuracy for local features was numerically higher than shape (*M*_shape_ = .83 vs. *M*_feature_ = .84). Together, these results suggest that shape may not be necessary to recognize familiar everyday objects, but that that local features are sufficient.

Next, we asked, how robust is the ability to use local features for familiar object recognition. To this end, we performed a follow-up experiment (*n* = 50) wherein we parametrically varied the stimulus duration (300, 136, 68, 34 ms; Fig. 1D). Here again we found that performance using local features was above chance (0.16) at all durations, even at the fasted speed of 34 ms (*ps* < .001, *ds* > 0.76, BF₁₀ > 100). This finding suggests that recognition using local features is robust and was not the result of inferential reasoning or top-down processing in the previous experiment. Moreover, there were no differences between shape and feature performance at 300 and 136 ms stimulus durations (*ps >* .234, *ds* < 0.17, BF_10_ < 0.28) and it was only at the two fastest speeds (68 and 34 ms) that shape performance surpassed local feature performance (*ps <* .001, *ds* > 0.82, BF_10_ > 100). Nevertheless, the finding that recognition using local features was above chance even at speeds faster than the duration of a typical visual fixation (55) or a saccade (56), suggests that our ability to use local features for everyday object recognition is robust. Together with our first experiment, these findings demonstrate that not only are local features sufficient for naturalistic, everyday object recognition, but humans’ abilities to use local features is robust even at extremely short durations.

### Shape supports one-shot learning

Our results showed that shape may not be necessary to recognize familiar everyday objects, and that local features are sufficient. However, this finding stands at odds with an extensive literature highlighting the importance of shape information across perception and cognition (3–6, 8–11). If shape is not necessary for everyday object recognition, then what exactly is the functional role of shape? One hypothesis is that shape may be especially useful when first learning new object categories. Whereas local features may vary considerably across exemplars of a category (12), shape remains remarkably stable (5, 11, 14, 57, 58). This stability may be especially important when initially learning a new category because a single exemplar provides sufficient information about the appearance of other category members.

To test whether shape supports rapid category learning, we presented participants with a stimulus set that was novel to participants but possessed a real-world category structure, namely instruments from different cultures. Participants (*n* = 50) were trained to label one exemplar per category and then tested with a held-out set of stimuli using the same procedure as before (Fig. 2A).

**Figure 2.**
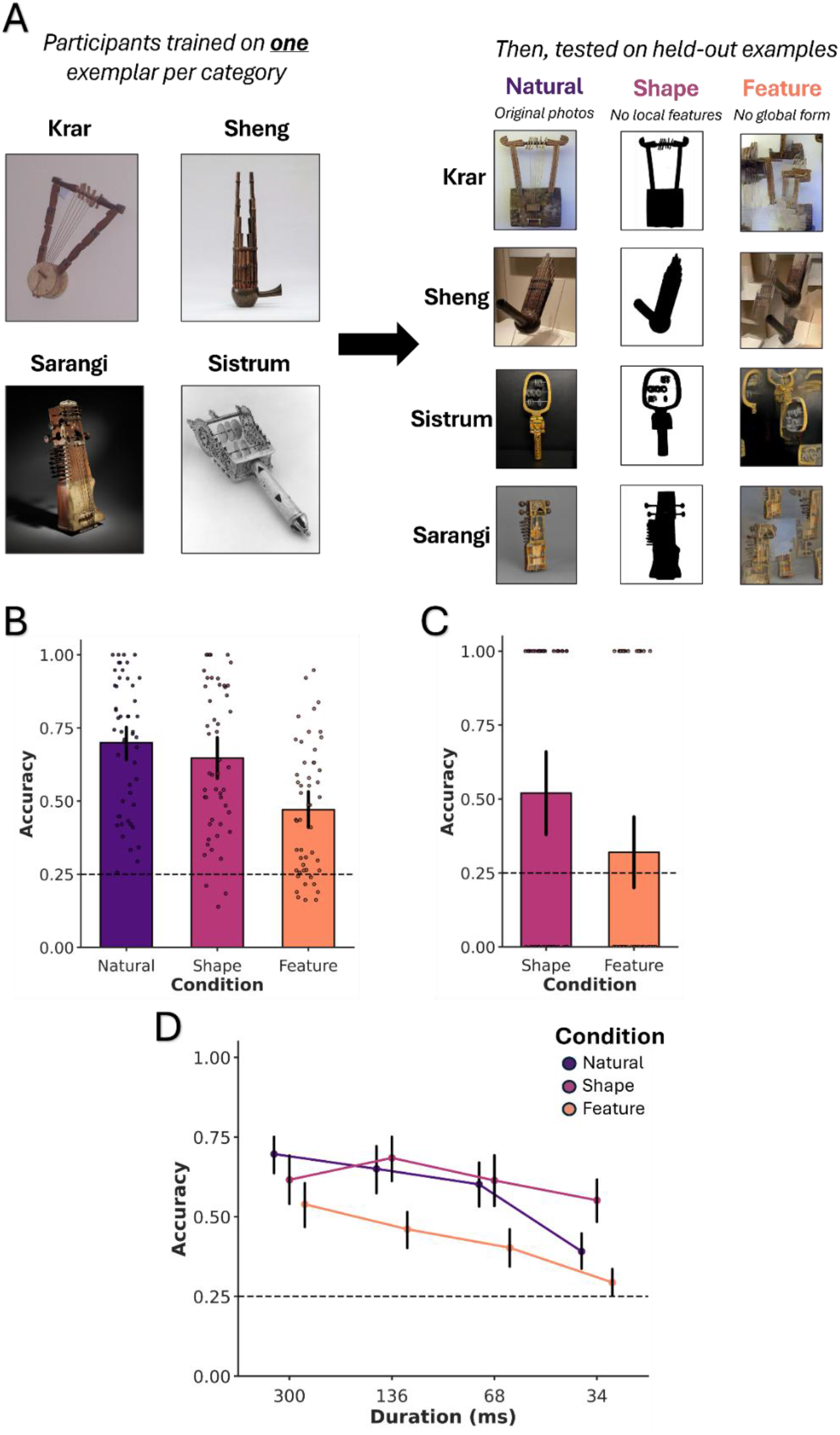
One-shot categorization stimuli, conditions, and results. (A) The stimulus set consisted of novel musical instruments that possess a real-world category structure. Participants were taught category labels for one exemplar from each instrument category, then tested on a held-out set of exemplar objects. (B-D) Participants performance for natural, shape, and feature conditions in the (B) consistent duration, (C) one-trial only, and (D) variable duration experiments. Individual points correspond to each participant’s mean accuracy on that condition or condition/stimulus duration. Error bars correspond to the upper and lower bounds of 95% confidence intervals. Dashed lines correspond to chance performance (0.25).

One-sample *t*-test comparisons to chance (0.25) revealed participants exhibited above chance performance for natural (*M* = .70), shape (*M* = .65), and local feature (*M* = .47) conditions (*ps* < .001, *ds* > 1.00, BF₁₀ > 100; Fig. 2B left). Next, we tested whether there were any differences between our conditions using a repeated-measures ANOVA with stimulus condition as the within-subjects factor. Our results revealed a main effect of condition on performance (*F*(2,98) = 46.36, *p* < .001, *n^2^_p_* = .49, BF_10_ > 100). Post-hoc multiple comparisons (Holm-Bonferroni corrected) revealed that performance on the natural condition was statistically higher than the shape condition (*t*(49) = 2.54, *p* = .014, *d* = 0.23, BF_10_ = 2.76), though the effect sizes and Bayes Factor suggest only weak or anecdotal evidence of a difference (59). Performance on the feature condition was also significantly lower than the natural (*t*(49) = 8.91, *p* < .001, *d* = 0.98, BF_10_ > 100) and shape conditions (*t*(49) = 6.34, *p* < .001, *d* = 0.76, BF_10_ > 100), suggesting that features are less informative for one-shot learning.

However, because the order of conditions was randomized for each participant and they completed multiple trials per condition, it is possible participants were actively learning about each object’s category structure during the test phase. From this perspective, performance on each condition may not reflect a one-shot learned category per se. To test this possibility, we carried out a follow-up experiment (*n* = 100) with an identical training regime and stimulus set, but participants were tested with only a single trial, thereby allowing us to evaluate their capacities without additional learning at test time. Participants were tested between-subjects in one of two conditions, shape *(n* = 50) or feature (*n* = 50) and performance was compared between each group (Fig. 2C).

Contrary to the results above, one-sample t-tests comparing both shape and feature conditions to chance (0.25) and found that performance on shape was above chance (*t*(49) = 3.78, *p* < .001, *d* = 0.54, BF_10_ > 100; Fig. 2C), whereas performance using local features was not (*t*(49) = 1.05, *p* = .299, *d* = 0.15, BF_10_ = 0.30). The post-hoc comparisons between shape and feature conditions revealed that performance on the shape condition was significantly higher than performance on feature condition (*t*(98) = 2.05, *p* = .043, *d* = 0.41). However, Bayes factor suggests only weak evidence in favor of the alternative hypothesis (BF_10_ = 1.33), likely due to the variability of participants only performing one trial. Nevertheless, these findings illustrate that shape, but not local features, plays a key role in one-shot category learning.

Finally, we asked, how robust is our ability to use shape for one-shot category learning. Mirroring our first set of experiments, we varied the stimulus durations (300, 136, 68, and 34 ms). As before, participants (*n* = 50) were trained on one exemplar and tested on a held-out set using the same procedure and conditions (Natural, Shape, Feature) as our previous experiment.

We found that performance on natural, shape, and feature conditions was above chance (0.25) at all durations (*ps* < .039, *ds* > 0.30, shape & natural : BF_10_ > 100, feature: BF_10_ > 2.65; Fig. 2D). Post-hoc multiple comparisons (Holm-Bonferroni corrected) between conditions showed that performance on natural condition was significantly greater than feature condition at all durations (*ps* < .008, *ds* > 0.41, BF_10_ > 6.49). By contrast, natural performance was only modestly higher than shape at 300 ms (*t*(49) = 2.33, *p* = .048, *d* = 0.34, BF_10_ = 1.97). There was no significant difference between performance on natural and shape conditions at 136 and 68 ms, (*ps* > .310, *ds* < 0.15, BF_10_ < 0.22), and shape performance was in fact higher than natural at 34 ms (*t*(49) = 4.32, *p* < .001, *d* = 0.68, BF_10_ > 100). Performance on shape was also higher than feature at all durations (*ps* < .048, *ds* > 0.33, BF_10_ > 1.17). Thus, although participants were able to identify objects equally well using local features and shape when objects were familiar, there was a strong advantage to using shape to categorize newly learned objects. Indeed, shape even matched or surpassed participants’ recognition of natural images in some cases, suggesting it may play a privileged role in rapid object learning. However, the fact that local feature performance was also above chance in many cases, even at fast durations, also illustrates the utility of using local features for object learning.

### Models rely on local features

Next, we explored how well human performance in our experiments matches that of visual DNNs, namely ConvNext and ViT (Fig. 3). These two models were chosen for their high-performance on visual recognition tasks, and because they were trained on transparent datasets, ensuring that our novel objects are unique to them as well. Like humans in the consistent duration experiment, models were evaluated using a 9-way classification procedure using a k-nearest neighbor classifier (KNN; see Methods).

**Figure 3.**
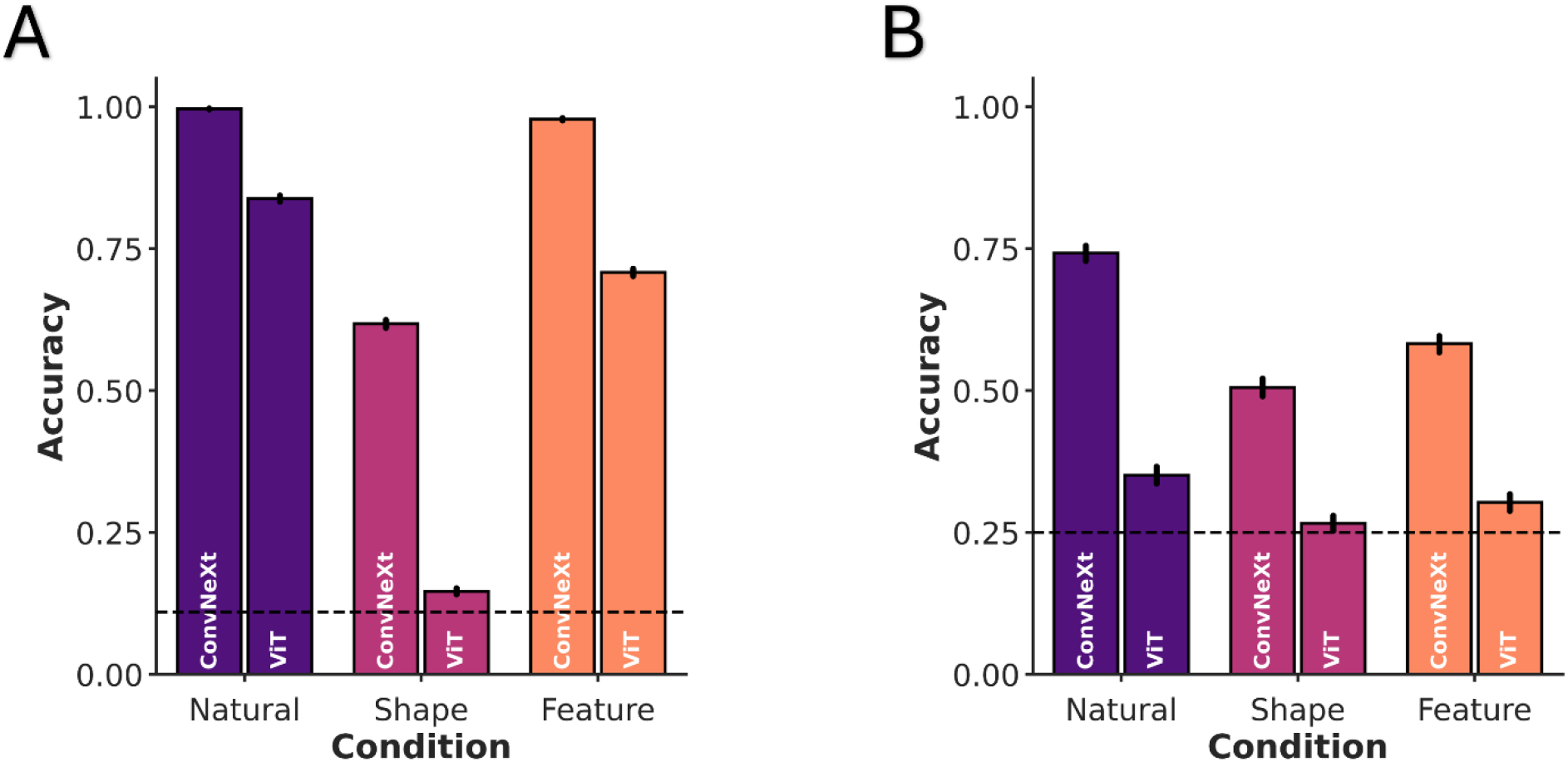
Model performance across experiments (A-B). Models performed above chance on all conditions for both the (A) familiar and (B) novel objects. (A-B) However, in all cases, performance was highest on feature condition compared to shape condition. Error bars correspond to the upper and lower bounds of bootstrapped (10,000 samples) 95% confidence intervals. Dashed lines correspond to chance performance (familiar: 0.11; novel: 0.25).

For familiar objects, we found that, like humans, both ConvNext and ViT exhibited the strongest performance on the natural condition (Fig. 3A; natural: *M*_ConvNext_ = .99, 95% CI [.995, .997]; *M*_ViT_ = .84, 95% CI [.83, .84]). And, consistent with prior work (16, 19, 21), DNNs exhibited significantly stronger performance on the local feature condition (*M*_ConvNext_ = .98, 95% CI [.976, .98]; *M*_ViT_ = .71, 95% CI [.70, .71]), than the shape condition (*M*_ConvNext_ = .62, 95% CI [.61, .62]; *M*_ViT_ = .15, 95% CI [.14, .15]), with both models performing more than 30 points higher using local features than shape. Thus, whereas humans exhibited equally high performance in both shape and feature conditions, DNN performance was substantially stronger in the feature condition.

For novel objects, we found that, like humans, models performed above chance (0.25) on all conditions. Furthermore, like humans in the consistent duration experiment, models performed best on the natural condition (Fig. 3B; natural: M_ConvNext_ = .74, 95% CI [.73, .76]; M_ViT_ = .35, 95% CI [.34, .37])). However, unlike humans, models performed better on the local feature condition (feature: M_ConvNext_ = .58, 95% CI [.57, .60]; M_ViT_ = .30, 95% CI [.29, .32]) than on the shape condition (shape: M_ConvNext_ = .51, 95% CI [.49, .52]; M_ViT_ = .27, 95% CI [.25, .28]), such that the lower bound of ViT’s CI overlapped with chance. Thus, whereas humans clearly prioritized shape information when categorizing novel objects, models relied on local features.

### Recognition using shape and local features is preserved across development

We found that, like DNNs, human adults could use local features to identify familiar everyday objects, but shape seems to be more crucial for one-shot learning. One explanation for adult participants’ remarkable performance on the local feature condition is that, like DNNs, they are able to draw on extensive visual experience to recognize objects. From this perspective, one might predict that populations who have less visual experience, such as children, might rely on shape more than local features across all tasks. To test this possibility, we repeated the familiar and novel categorization experiments with five-year old children (*N* = 100). The design was identical except participants performed a two-way classification for familiar everyday objects and a three-way classification for novel and recognition experiment (see Methods).

For familiar object recognition, we found that, like adults, children’s performance on all conditions was above chance (0.50) (*ps* < .001, *ds* > 2.20, BF_10_ > 100; Fig. 4A). Post-hoc comparisons (Holm-Bonferroni corrected) between performance on shape (*M* = .81) and feature (*M* = .81) conditions revealed no significant differences (*t*(49) = 0.06, *p* = .949, *d* = 0.01, BF_10_ = 0.15). These results suggest that by at least 5-years-of-age, children can readily use local features for everyday object recognition and shape may not be necessary.

**Figure 4.**
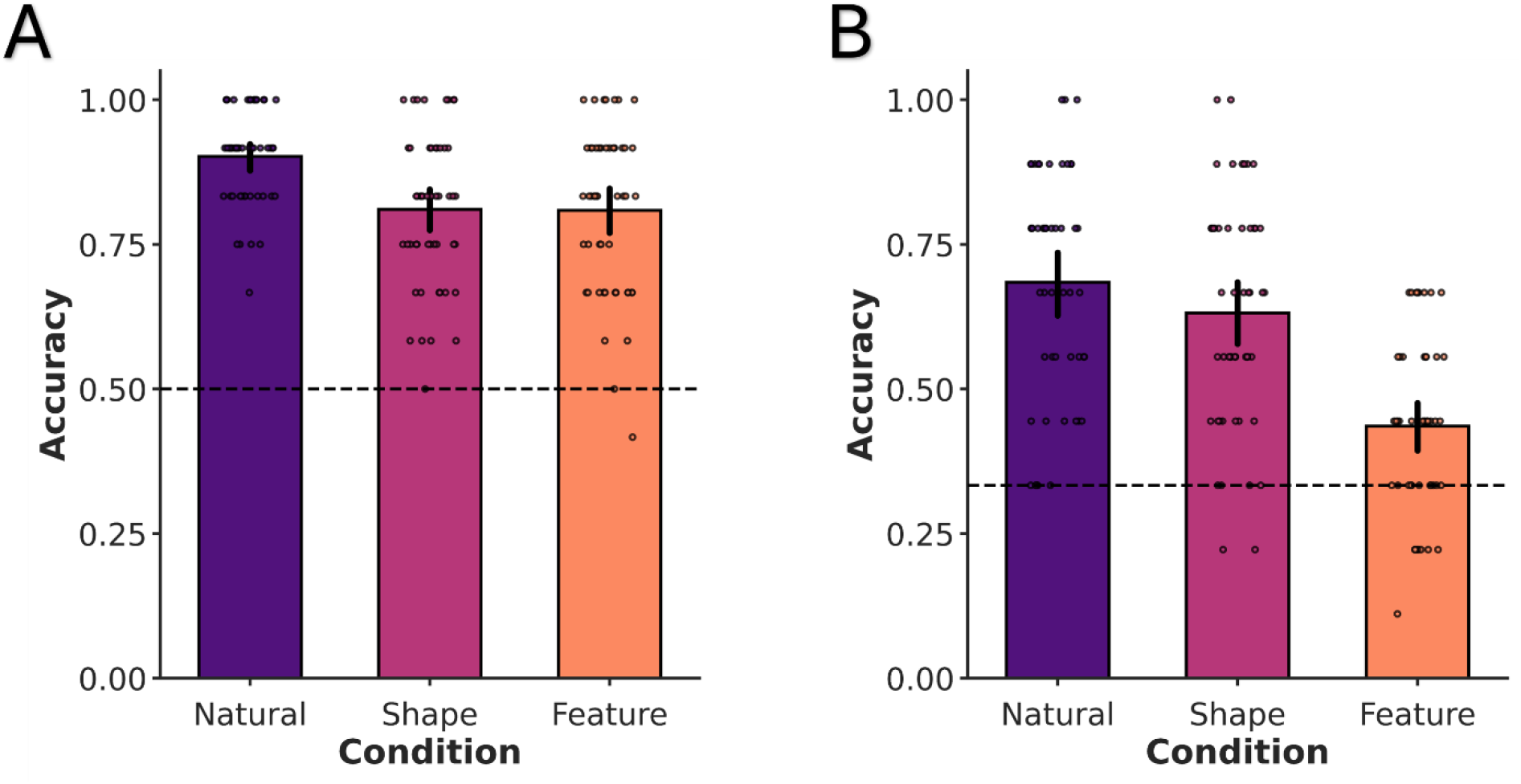
Developmental stability in object recognition and learning. (A-B) Children’s performance for natural, shape, and feature conditions in the (A) familiar, object recognition experiment and (B) novel one-shot learning experiment. (A-B) Participants’ performance on each condition was above chance in both experiments. Individual points correspond to each participant’s mean accuracy on that condition. Error bars to the upper and lower bounds of 95% confidence intervals. Dashed lines correspond to chance performance (familiar: 0.5; novel: 0.33).

For the one-shot categorization task, we found again, like adults, children’s performance on all conditions was above chance (*ps* < .001, *ds* > 0.71, BF_10_ > 100; Fig. 4B). Post-hoc comparisons (Holm-Bonferroni corrected) revealed that, like adults, children’s performance on the shape condition (*M* = .63) was significantly higher than performance on the feature condition (*M* = .44) (*t*(49) = 7.04, *p* < .001, *d* = 1.07, BF_10_ > 100). These results suggest that using shape for one-shot categorization is privileged even early in development. However, it is important to note that even children could use local features to some extent here as indicated by their above chance performance on feature condition. Altogether, these findings illustrate that there is developmental stability in how children use shape and local features to recognize and learn objects.

## Discussion

Although decades of research have emphasized the importance of shape for perception and cognition, the current best computational models of human vision do not prioritize shape, but instead rely on local visual features. Thus, in the current study, we reexamined the importance of shape and local features in human object recognition and one-shot learning. We found that local features, in the absence of global shape information, were sufficient for adults and children to recognize familiar objects. Recognition using local features was remarkably robust, with accuracies often matching those in the shape condition, and remaining high even at stimulus speeds as fast as 34 ms. These findings suggest that, like DNNs, humans can use local features to identify common objects and that shape may not be as important for everyday object recognition as previously believed. Moreover, these findings may also explain how DNNs often match human visual processing without representing shape. Yet, although shape may not be necessary for recognizing familiar objects, we found it may play a privileged role in category learning, allowing both children and adults to categorize novel objects after viewing just one example. Altogether, these findings reconceptualize our understanding of the role of shape in human visual perception and suggest that local features are more important than previously considered. These results also demonstrate that despite not using shape, DNNs still capture a fundamental aspect of human vision and perception. Finally, while shape may not be necessary for everyday object recognition, it serves as a vital cue in supporting rapid category learning.

### Limitations and alternative explanations

Given participants’ performance in using local features to recognize familiar objects, it is important to consider alternative explanations. For instance, one explanation for participants’ strong performance using features is that participants succeeded on this task by relying on top-down inferential processes that let them deduce the object category. This possibility, however, is not likely because, at their slowest duration, stimuli were presented for only 300 ms and were backward masked—a standard procedure for reducing recurrent and top-down visual processing (41, 60, 61). Moreover, accuracy on the feature condition remained above chance at extremely fast stimulus durations, even for novel objects. The finding that participants still performed accurately at these speeds suggests recognition using local features occurs rapidly via a feedforward pass through the visual system. Together, this suggests that participants’ strong performance using local features is not the result of top-down inferential reasoning.

However, another possible concern is that participants succeeded using local features because this manipulation did not sufficiently scramble the images and retained large diagnostic object properties. Although there is variability in the appearance of local features, with some almost entirely texture-like and other retaining object-like elements (Fig. 1A), all stimuli were algorithmically generated from outputs of a high-level DNN layer using the same procedure (see Methods). This approach allowed us to directly test our main questions: how well humans (1) recognize objects whose global shape is disrupted via spatial scrambling, and (2) whether humans are able to use the same visual cues as DNNs. Importantly, because both the local feature and shape conditions were algorithmically generated for the same set of images, this further minimizes the possibility of experimenter bias in selecting stimuli that benefit one set of images over the other. Indeed, algorithmically transforming images into shape silhouettes can also create unintuitive object viewpoints (Fig. 1A), and removed diagnostic information. Thus, our stimulus generation procedure was applied to all images in an unbiased manner based on the specific hypotheses laid out by feature and shape-based theories (30, 33).

Nevertheless, it is an open question for what exactly defines a local feature (62), and how minimal, or texture-like, a feature can be while still supporting recognition. An interesting possibility is that features of all scales contribute meaningfully to recognition. For instance, prior work has demonstrated that even unrecognizable texforms contain valuable category-level information about animacy and size (63–65). Future studies could isolate different types of local features and systematically test whether certain types might be more diagnostic than others.

In contrast to familiar object recognition, we found that shape was more important than features for one-shot object learning. However, here we used a much smaller stimulus set where all objects were from the same superordinate category (i.e., instruments). Is it possible that the local features of each novel object class were more similar to one another and therefore less informative and distinguishable? This possibility also seems unlikely. First, the familiar recognition experiments also contained object labels that were within the same superordinate-level category (e.g., animate, inanimate natural, and inanimate artificial). Second, DNNs showed stronger recognition performance using local features than shape in novel object learning experiments, suggesting that feature stimuli still possessed meaningful visual information that could be extracted from the image despite the disruption of global form.

### Reconceptualizing the role of shape and local features in object recognition

Historically, shape has been considered to be the principal cue underlying object recognition (4–6, 11, 20, 66), particularly for basic-level categories (13, 14). Local features by contrast, were, at best, treated as a secondary cue that may support subordinate-level classification (5, 67, 68). However, our results challenge this characterization and demonstrate that local features can readily support basic-level object recognition, and shape may not even be necessary in this context. Indeed, performance using local features alone often matched performance using shape and retained high accuracy at stimulus durations that were shorter than a visual fixation (55). This suggests that when viewing objects in our typical visual environment, which is largely comprised of familiar objects and rarely involves strong time constraints, shape may not be needed to solve visual recognition tasks. Instead, like DNNs, humans may use local features to quickly identify the objects in their environment.

These findings are consistent with a growing body of evidence that suggests the ventral visual pathway, the primary neural substrate underlying object recognition, may also primarily rely on local features (30). Indeed, like DNNs, individual neurons in the ventral pathway, show stronger tuning to local object features than complete object forms (48, 69, 70). Similarly, at the neural population level, the multivariate and univariate responses of the ventral pathway show similar responses to intact and spatially scrambled objects that preserve high-level features (33, 65). This local feature coding principle is not unique to objects but even extends to categories such as faces, that were historically thought to be defined by holistic properties (71–73). Our work builds upon these findings by showing that the local feature coding properties of the ventral pathway are also behaviorally relevant, allowing humans to quickly recognize objects in their environment, and thereby explain how DNNs achieve strong fits to the ventral pathway without clear shape representations.

### Shape supports rapid object learning

Although shape may not be necessary to recognize familiar objects, our findings suggest that it is crucial for rapid object learning. Whereas local features can be highly variable across exemplars within the same basic-level category (5, 14, 66, 67), and therefore individuals may require more visual experience to learn the distribution of possible features for a category, global shape information remains stable across category members. A reliance on local features may explain why DNNs require much more visual experience than humans to learn new object concepts (11, 20), as well as why the ventral pathway has a protracted, experience-dependent developmental trajectory (74, 75). By contrast shape is remarkably stable across exemplars of a category (5, 11, 57, 58), which explains why it forms the basis of many basic-level object categories (13, 14) . This stability means that individuals can, after seeing just one category example, accurately infer the appearance of other category members, making it a privileged cue for rapid category learning.

However, given this framework, one might have predicted that young children, who have much less visual experience than adults, would have been worse at using local features than shape to recognize objects. However, our results with 5-year-old children showed nearly an identical pattern to adults, wherein features were sufficient and there was no significant difference between feature and shape performance. These findings suggest that by at least age five, children have gained enough visual experience to learn the distribution of local features of common categories. Future work may examine when such local feature representations arise, and test different age groups to determine whether younger age groups prioritize shape more than features. It is important to also note that both adults and children were often able to use local features in the one-shot learning experiments, even at fast stimulus durations, though performance here was worse than shape. This finding illustrates that even if local features are not as stable of a cue as shape, they nevertheless exhibit sufficient stability to support rapid recognition.

### Models capture an important aspect of human visual perception

A long-standing mystery in computational neuroscience is how visual DNNs could provide such a strong neural match to the ventral visual pathway (36, 43–45, 49, 50), while neglecting an important visual cue such as shape (16, 19, 21, 22). Our results provide a possible solution to this puzzle by showing that humans similarly do not need shape information when identifying familiar objects. And, as mentioned previously, this finding coincides with accumulating evidence that the ventral pathway may not represent global shape either, but is best described as a basis set of local features (30, 33, 48, 65, 76). Thus, DNNs match to the ventral pathway is not simply an accidental byproduct of the fitting function but instead reflects a biologically meaningful visual process.

However, unlike DNNs, humans can accurately use shape information, in the absence of features, to recognize objects, and prioritize shape when shape and local features are placed in conflict with one another (19). There are several developmental constraints that may help the human visual system prioritize shape when first learning new objects. For instance, poor visual acuity during infancy may make local features blurry, thereby forcing infants to prioritize global visual properties such as shape (77, 78). Indeed, training DNNs on an infant-like blurry-to-clear visual sequence results in improvements to their visual recognition capacities across a range of domains (79–83). Additionally, infants appear to self-select object viewpoints that maximize shape information (84, 85), and training DNNs on headcam data from infants leads to improvements in the robustness and efficiency of learning (86–88). Thus, the human visual system appears to have adapted to use shape early in development when most objects are still novel and visual experience is minimal. Implementing DNNs with developmentally plausible mechanisms may therefore result in more human-like visual representations (89).

In addition to developmental constraints, the neural systems involved in shape perception may extend beyond the ventral visual pathway (30, 31, 90–96). Specifically, global shape is defined by the spatial structure of the object, that is, how object parts are spatially arranged, or configured, relative to one another (3, 30, 31, 97). Accumulating evidence suggests that such a representation may be computed by spatial processing areas of the dorsal visual pathway (31, 90, 91). Indeed, the dorsal pathway represents spatial configuration of features for both objects (30, 31, 90) and faces (73) and transmits this information to the ventral pathway to support categorization (30, 31, 92, 98). Disrupting parietal areas of the dorsal pathway (via TMS or damage) impairs participants’ ability to perceive global, but not local, object properties (73, 99, 100), and even reduces responses of the ventral pathway (101). Interestingly, the dorsal pathway appears to mature earlier than the ventral pathway in both humans (102–105) and monkeys (106–108), thereby providing a mechanism by which shape may support object learning early in development. Thus, integrative modeling of both dorsal and ventral pathways (109–112), may lead to more robust human like object representations and learning capacities.

## Conclusion

In conclusion, these findings reconceptualize our understanding of the role of shape and local features in human visual perception. We show that local features may be more important than previously considered, whereas shape may be less important for everyday object recognition tasks. Instead, shape may play an especially important role in novel object learning. These results disentangle longstanding debates regarding the biological match of DNNs to the human visual system and illustrate a human-specific mechanism for rapid visual learning.

## Methods

### Participants

Sample sizes, testing procedures, and primary hypotheses for adults and children were preregistered. Based on power analyses using pilot data, we sought to recruit 50 participants for each experiment. This resulted in a total of 300 adult participants and 100 child participants. Adults were recruited from Prolific and children were recruited through Children Helping Science.

Participants and individual trials were excluded based on three exclusion criteria. Trials were excluded if participants’ response time (RT) for that trial was faster than 100 milliseconds (ms) or 2 standard deviations greater than or less than the mean RT for that participant. Trials were also excluded if participants did not respond within the allotted trial response time (5 s). Finally, we excluded participants whose accuracy on the Natural condition was at chance or 2 SD below the mean accuracy of all participants in the natural condition. For children, an independent coder blind to the conditions of the study watched recordings of each child session and excluded any participants whose parents helped them or the child was off-task. Any excluded participants were replaced until the targeted sample size was reached. On the basis of these criteria, a total of 15 adults and 16 children were excluded from analyses.

Links to pre-registration for each experiment are below:

> Adult familiar object recognition consistent duration: https://aspredicted.org/e923jv.pdf
>
> Adult familiar object recognition variable duration: https://aspredicted.org/5vr4fp.pdf
>
> Adult novel object recognition consistent duration: https://aspredicted.org/py7376.pdf
>
> Adult one-shot object recognition variable duration: https://aspredicted.org/ch8fb3.pdf
>
> Child familiar object recognition: https://aspredicted.org/ar8xd3.pdf
>
> Child novel object recognition: https://aspredicted.org/58wf6n.pdf

### Stimuli

The stimulus set used in all experiments consisted of naturalistic images of real-world objects (Natural), shape silhouettes with disrupted local features (Shape), and scrambled images wherein the global form had been disrupted (Feature) (Fig. 1A & 2A). We transformed naturalistic images using two image synthesis algorithms to create our shape (113–115) and feature stimuli (33).

For shape stimuli, we generated 2D silhouettes using a combination of Grounding DINO (115) and the Segment Anything Model 2.1 (SAM 2.1) (113). For each image, a text prompt corresponding to the target object’s category label (e.g., “bear”) was passed to Grounding DINO, which returned a bounding box around the matching object (box threshold = 0.4, text threshold = 0.3). This bounding box was then provided as a prompt to SAM 2.1, whose transformer-based mask decoder identified the pixels belonging to the object within the box. This produced a pixel-wise probability mask, which was subsequently thresholded to produce a binary segmentation mask thereby creating a 2D silhouette of the target object where the foreground pixels represented the segmented object and background pixels represented non-object regions. The final generated 2D segmentation masks were used as the shape stimuli. For local feature stimuli, we employed the texture synthesis algorithm from Jagadeesh and Gardner (33). This algorithm uses VGG19, an ImageNet-trained convolutional neural network, to produce spatially scrambled images that preserve high-level visual features. Feature maps were extracted from ‘pool4’ of VGG19 and were scrambled using a 1 × 1 spatial constraint, which allowed features to be scrambled across the entire image (for additional details on the image synthesis algorithm see (33)).

Familiar object stimuli were comprised of basic-level categories selected from the EcoSet image database (116). We selected an equal number of animate, inanimate natural, inanimate artificial objects. In total there were 18 categories for adults and 12 for children, with each category comprised of 10 exemplars each. The stimuli in the one-shot learning experiments were comprised of four instruments from different cultures around the world, namely a krar, sistrum, sheng, and sarangi (10 exemplars each). An additional three exemplars of each musical instrument were used for the training phase of experiments, with only one of the three training exemplars randomly selected for each participant. To simplify the task for children, we used three instruments, specifically a sarangi, sheng, and sistrum (5 exemplars each) and they conducted a three-way classification rather than a four-way classification like in the adult version. We selected these instruments because they possess a real-world category structure, comprised of visually distinct exemplars, but are unfamiliar to American participants.

### Human Testing Procedure

We adapted an object recognition paradigm from Wichman et al. (60) to test the degree to which shape or local features are necessary or sufficient for object recognition in adult humans. Participants were presented with common every-day object images in one of three conditions: (1) a ‘natural condition’ where the image is shown in its original undisrupted state, (2) a ‘shape condition’ which removes the local features but preserves the global shape information by presenting objects as silhouettes, and a (3) ‘feature condition’ which disrupts shape information, but preserves local features by spatially scrambling the image (Fig. 1A).

For all experiments, the basic structure of the task was the same (Fig. 1B). Each task began with instructions followed by a practice block to ensure participants understood the task. For familiar object recognition tasks, this involved a practice set of objects separate from the test object classes. For the novel learning task this practice also served to teach participants the novel object labels using one exemplar per object class. Here, participants were randomly assigned one image example from each novel object category.

Stimuli in the practice block were presented for 1 s with no time limit for responses, and feedback was given as a green check mark for correct and a red ‘X’ for incorrect responses. In the test phase with a consistent duration, trials began with a fixation cross in the center of the screen (500 ms), followed by the object image (300 ms), a mask (200 ms), and then a gray screen with numbered labels (5 s) corresponding to different object categories (e.g., 1. dog, 2. cat, 3. Airplane etc.; Fig. 1B). In the variable duration experiments, the stimulus speed varied from 300 ms to 34 ms (300, 136, 68, & 34 ms) and progressively decreased as the experimental block progressed.

For each task participants were asked to press the number on their keyboard corresponding to the object label. If participant did not respond within the allotted time, their responses were marked as “no response,” and a screen appeared (2 s) prompting them to respond more quickly in the next trial.

For familiar object recognition task, there were a total of 9 labels presented on each trial out of 18 possible object class labels (in a random arrangement). Response labels corresponded to each of our superordinate object categories, with three object classes corresponding to the same superordinate category (e.g., 3 animate, 3 inanimate natural, and 3 inanimate artificial). Because the variable duration versions of our experiments were potentially more difficult, we limited the number of labels to 6, with 2 from same superordinate category. For novel object experiments, as there were only four object classes in total, the labels were identical for each test trial, but in a random arrangement.

Participants completed three blocks of test trials, each consisting of one experimental condition (natural, feature, condition), and the block order was randomly assigned to each participant. In the adult familiar version of the experiment, participants completed 72 trials for a total of 216 trials before trial exclusion. In the novel version, participants completed 40 trials per experimental block for a total of 120 trials before trial exclusion.

For the child versions of each experiment, the task structure and design were the same, with a few modifications to ensure the task was accessible for children. Like adults, children were required to recognize a rapidly presented object image (300 ms) that was backward masked (200 ms). However, there was no time limit to respond and instead of performing a 9-way classification in the familiar object recognition experiment, children performed a 2-way classification. For our one-shot learning experiment, instead of performing a 4-way classification, they performed a 3-way classification. Furthermore, the number of trials per condition block was reduced to 12 trials per block for both familiar and novel. Children responded by saying or pointing at the label and then the parent would press the corresponding key.

### Model Testing Procedures

We tested two DNNs: ConvNext and ViT. These models were selected because of their high performance on visual recognition tasks (40–42) and because they were trained on transparent dataset (i.e., ImageNet), thereby ensuring that they were not already exposed to the novel objects or stimulus conditions of the current study. In contrast, other high performing vision models (e.g., CLIP, DINOv2) are trained on massive opaque datasets from the internet, making it impossible to know if they were already exposed to the stimuli used here.

We tested model recognition accuracy using the same stimuli and a comparable procedure as adults. In the familiar recognition task, adults were required to determine which of 9 semantic labels (e.g., dog, cat, airplane etc.) corresponded to the target image. To accomplish this task, participants must be able to match the stored object representation associated with each label (e.g., previously seen dogs) with the stimulus they saw in the trial. To approximate this process, we trained a K-Nearest Neighbor (KNN) classifier, using the feature activations from the penultimate layer of each model on images from the training set of Ecoset, which did not include the images adults were tested with (c116). Images consisted of naturalistic objects humans see every day and were from the same categories adults were tested on. For each stimulus category, we provided 300 images for classifier training (5,400 images total). To emulate the 9-way classification task adults performed, on each evaluation trial we trained the classifier on feature activations training images corresponding to the target object, as well as training images from 8 other classes, with the constraint that there were a total of 3 classes from each superordinate category like in the adult experiment. All 18 stimulus categories from the adult experiment were tested, and each exemplar (10 exemplars per category) was tested 100 times on each condition with a randomly selected subset of labels.

For the novel object recognition task, we trained a KNN classifier on the feature activations from the penultimate layer of each model on a single exemplar from each category (same exemplar adults were trained with) and then tested them on the same held-out images adults were tested with. Model recognition accuracy was evaluated on each condition in a 4-way classification on 10 images per stimulus category and each image was repeated 100 times.

### Statistical Analyses

For each version of the experiment, performance on each condition was compared to its respective chance level using one-sample *t*-tests. Next, one-way repeated measures were conducted, and planned post-hoc comparisons were carried out between conditions. However, for one-trial only experiment, a between-subjects design was employed to compare participants who completed the feature condition vs. those who completed the shape condition. For all analyses, both inferential and Bayesian statistics were run which allowed us to detect the effect (Inferential) and how much evidence we had for or against the hypotheses (Bayesian). These two analyses showed qualitatively similar results in all cases. Furthermore, because some datasets were not normally distributed, all data was analyzed with both parametric and non-parametric analyses. The results of the parametric and non-parametric statistics did not qualitatively differ and showed similar effect sizes in all cases.

